# Frequency of five *Escherichia Coli* pathotypes in Iranian adults and children with acute diarrhea

**DOI:** 10.1101/725952

**Authors:** Sana Eybpoosh, Saeid Mostaan, Mohammad Mehdi Gouya, Hossein Masoumi-Asl, Parviz Owlia, Babak Eshrati, Mohammad Reza Montazer Razavi Khorasan, Saeid Bouzari

**Author notes:** **Corresponding author** E-mail address (SB). These authors contributed equally to this work (Sana Eybpoosh, Saeid Mostaan).

## Abstract

**Background:** knowledge about the distribution of *Escherichia Coli* (*E. coli*) pathotypes in Iran is limited to studies with small scale and limited scope. This nation-wide survey aims to provide a more generalizable estimate of pathogenic *E. coli* distribution in Iran.

**Methods:** During January 2013 and January 2014, stool samples were collected from 1306 acute diarrhea cases of 15 provinces. Culture-positive *E. coli* samples were tested with PCR for detection of five *E. coli* pathotypes (STEC, ETEC, EPEC, EAEC, and EIEC). Frequency of these pathotypes was estimated for different provinces, age groups, and months/seasons.

**Results:** Of 1305 diarrheal samples, 979 were *E. coli*-positive (prevalence: 75.0%; 95% CI: 72.6, 77.3%). Pathogenic *E. coli* was detected in 659 diarrheal samples (prevalence: 50.5%; 95% CI: 47.8, 53.2%). STEC and EIEC was the most and the least frequent pathotypes (35.4% and 0.3%, respectively). ETEC (14.0%) and EPEC (13.1%) were the second and the third frequent pathotypes, respectively. EAEC was not highly prevalent (4.3%). Fars and Razavi Khorasan provinces had the highest and lowest frequencies (88.7% and 34.8%, respectively). *E. coli* pathotypes were more frequent in warmer (i.e., spring and summer) than cooler (i.e., fall and winter) seasons. The highest frequency of pathogenic *E. coli* was observed in infants and children under 5 years (73% each). There was no association between sex and pathogenic *E. coli* infection.

**Conclusions:** Diarrheagenic *E. coli* may be an important cause of acute diarrhea in adults and children in Iran. STEC and ETEC seem to be widespread and show a peak in warmer seasons. This finding could impact the recommended use of STEC and ETEC vaccines during warmer seasons, especially for infants, young children and elderlies. Monitoring the rate of diarrheagenic *E. coli* infection, *E. coli* serotypes, and their antibiotic resistance is recommended for evaluations of time-trends and effectiveness of interventions.

**Author summary:** *Escherichia coli*, also known as *E. coli* is a bacterium of the genus *Escherichia* that is normally found in the lower intestine of human. Most *E. coli* strains are harmless, but some can cause infection in the gastrointestinal tract, causing diarrhea. These pathogenic *E. coli* strains are classified based on their mechanism of pathogenesis. In this regard, five important *E. coli* strains include Shiga toxin-producing *E. coli* (STEC), enterotoxigenic *E. coli* (ETEC), enteropathogenic *E. coli* (EPEC), enteroaggregative *E. coli* (EAEC), and enteroinvasive *E. coli* (EIEC). In a national survey conducted in Jan 2013 till Jan 2014, we collected 1305 diarrheal samples from 15 (out of 31) provinces of Iran. Of these, 979 samples (75%) were *E. coli*-positive in the culture test. Molecular tests showed that 659 samples were pathogenic *E. coli*, suggesting that 50.5% of the diarrhea cases were induced due to pathogenic *E. coli* infection. The most prevalent pathogenic *E. coli* strains in Iran were STEC (35.4%) and ETEC (0.3%), and were more commonly detected in warmer seasons, infants, and children less than five years. So, the use of vaccines, especially for STEC and ETEC, during warmer seasons and for infants, young children and elderlies are recommended.

## Introduction

Diarrheal diseases are among major cause of illness and death, especially among children (1). Deaths are mainly caused by loss of water and electrolytes resulting from intestinal malabsorption or increased secretion (2). Although the condition is considered as a major health concern worldwide, its prevalence and adverse effects are more prominent in lower income and less developed countries. Projections have estimated that by 2030, diarrheal diseases remain as one the top 10 causes of death in low income countries (1).

*Escherichia coli* (*E. coli*) is a gram-negative bacterium found naturally in the intestinal tract of human. A subset of *E. coli* is capable of causing diarrheal disease (known as diarrheagenic *E. Coli* [DEC]), with some of its strains leading to severe diarrhea and even death. Different DEC strains (known as pathotypes) have been determined relatively recently, including Shiga toxin–producing *E. coli* (STEC), enteropathogenic *E. coli* (EPEC), enterohemorrhagic *E. coli* (EHEC), enterotoxigenic *E. coli* (ETEC), enteroaggregative *E. coli* (EAEC), enteroinvasive *E. coli* (EIEC) and diffusely adherent *E. coli* (DAEC) (3).

The main virulence factor of STEC is a phage-encoded potent cytotoxin. The cell toxicity effect of this pathotype was also demonstrated on Vero cells, resulting in a parallel nomenclature of Shiga/Vero toxin-producing *E.coli* (STEC) and VTEC, respectively (4). ETEC is characterized by its ability to elaborate the heat-stable and/or heat-labile enterotoxins, and is noted to be the major cause of travelers’ diarrhea globally (5). Like many strains of STEC, EPEC harbors the *Eae* gene, encoding the outer membrane protein “intimin”. This enables both pathotypes to attach epithelial cells of the intestinal tract (6). STEC are distinguished from EPEC strains by harboring the Shiga toxin–encoding gene. While EPEC strains lack chromosomal Shiga toxin-encoding genes, they harbor the EPEC adherence factor (EAF) virulence plasmid, which encodes the *Bfp* fimbriae. EAEC is recently defined as a DEC, and is increasingly reported in lower income countries (7). Recent identification of virulence factors under the control of the AggR regulator have categorized EAEC among diarrheal pathogens (8). EIEC are closely related to *Shigella species*, in terms of their biochemical and physiological properties and their genetic structure. Both *Shigella* and EIEC encode common putative virulence genes and secret toxins that allow these pathogens to invade intestinal tract mucosa, move within epithelial cells, and penetrate neighboring cells (9, 10).

Existing evidence suggest a location-specific distribution of *E. coli* pathotypes. For example, some of the serotypes of STEC, including O157:H7, are well recognized in the United States and Canada as the major cause of hemolytic uremic syndrome. The non-O157:H7, however, is prevalent in Latin America, Australia, and Europe. ETEC is characterized as a major cause of travelers’ diarrhea, in almost all parts of the world (5). In Iran, very limited data is available about the existence and distribution of DEC in human. This may be the result of the paucity of large-scale and well-designed systematic epidemiological studies, and the absence of a surveillance system (11). For example, many of the available studies have focused on infants and children under 5 years of age (12–17). The very few studies conducted on adolescents and adults are limited to small samples, few geographical areas, and limited *E. Coli* pathotypes (18–21). The sampling procedure in all studies available from Iran is non-probabilistic. The sampling location of most available studies is also limited to one city. These factors limit the generalizability of the findings of existing studies.

Given current knowledge gap, we performed this nationally-representative survey to elucidate the role of five *E. Coli* pathotypes in the epidemiology of diarrheal diseases, and to provide more recent estimates of the prevalence of DEC pathotypes among adults and children of Iran. It is hoped that the results of this survey be used for policy making for diarrhea management in Iranian population and provide a basic insight for vaccine design.

## Material and Method

### Ethics statement

This study was approved by the ethics committee in Pasteur Institute of Iran. All participants were informed about study objectives, assured about the confidentiality of their information, and gave their written informed consent to be included. For collecting sample from children and infants with diarrhea, a parent or guardian of any child participant provided informed consent on their behalf.

### Study design

A cross-sectional survey was conducted from January1, 2013 to January 30, 2014 to determine the prevalence and populations of *E. coli* in cases with acute diarrhea.

### Study population

The study population consisted of all Iranian nationals who were residents of target provinces of Iran and referred to the primary health centers with a chief complain of acute diarrhea (with or without bleeding). Acute diarrhea was defined as the acute onset of liquid and watery bowel movements more than 2-3 times a day. In order to identify the real distribution of *E. coli* in selected provinces, we excluded patients who reported to travel out of the province in the past two weeks. We also excluded diarrhea cases that were under chemotherapy, antibiotic therapy, or corticosteroid therapy as well as all immunologically compromised patients. Patients with chronic diarrhea were not included either.

### Sampling procedure

Of 31 provinces of Iran, 15 provinces were selected (sampling fraction= 50%). Provinces were sampled in a way to provide an even distribution of all geographic areas across the country. In each of the provinces three cities were selected. This resulted in the selection of 45 cities across the country. In each city, stool sample from diarrhea cases were collected at the second half of each month, from which a random sample were cultured for *E. coli* detection. *E. coli*-positive samples identified in culture method were then sent to the National *E. Coli* Reference Laboratory (NECRL) in Pasteur Institute of Iran (PII) for molecular analyses. Collecting samples from diarrhea cases that were occurred at the second half of each month was to homogenize the sampling time in all cities. Finally, a total of 1305 diarrhea samples were collected over a period of one year.

To ensure data quality and standardize data collection and classification in all field centers, we held a series of workshops in which participating staffs were thoroughly trained about the procedures under their responsibility. Trainings involved how to collect stool samples, do the cultures, interpret and record culture results, and transfer the samples to NECRL.

### Isolation and Identification of *E. coli*

#### Culture

Stool samples were cultured in the laboratories of the field centers. Samples that were confirmed as *E. coli-*positive, were transferred to NECRL for molecular analysis. For the culture test, stool samples were inoculated on MacConkey agar (Merck, Catalog No. 105465) at 37 °C for 24 hours. After O/N incubation (incubator: Vision Korea), results were checked, and five typical colonies on MacConkey agar (pink color) were selected and transferred to triple sugar iron agar media (Merck, Catalog No. 104728). Strains with *E. coli* characteristics were selected and transferred to SIM medium (Merck, Catalog No. 105470), Simmons citrate (Merck, Catalog No.102501) and MRVP broth (Merck, Catalog No. 105712) to check for the presence of *E. coli* characteristics. Indol-positive (Kovacs, Merck, Catalog No. 109293), MR-positive (Methyl red, Merck, Catalog No. 106076), VP-negative (KOH, SIGMA USA, Catalog No. P5958 and alpha naphthol-1 Catalog No. 70480) and citrate-negative samples were identified as *E. coli*. Identified *E. coli-*positive samples were incubated on LB (Merck, Catalog No. 110285). After O/N incubation, samples were centrifuged (Eppendorf Germany) and the final product was stored at −70 °C until used (22).

#### PCR

Eight *E. coli* virulence genes indicative of five *E. coli* pathotypes were selected and checked for using PCR test (Table 1). These included genes coding toxins (Shiga toxins [*Stxs*] *Stx1* and *Stx2*, heat-labile toxin [*LT*], and human heat-stable toxin [*ST*]), adhesion factors (*Eae* and *Bfp*), and invasion genes (*invE*). The primers used for probe amplification were either chosen from existing literature on virulence gene detection or designed from available gene sequences. Target genes in this study, were used to identify five *E. coli* pathotypes, including: (i) STEC, identified by *Eae*, as well as *Stx1* or *Stx2* or both; (ii) ETEC, identified by *LT* and *ST* genes; (iii) EPEC, identified by *Eae* and *Bfp* genes; (iv) EAEC, identified by *AA* gene; and (v) EIEC, identified by *invE* gene.

**Table 1.**
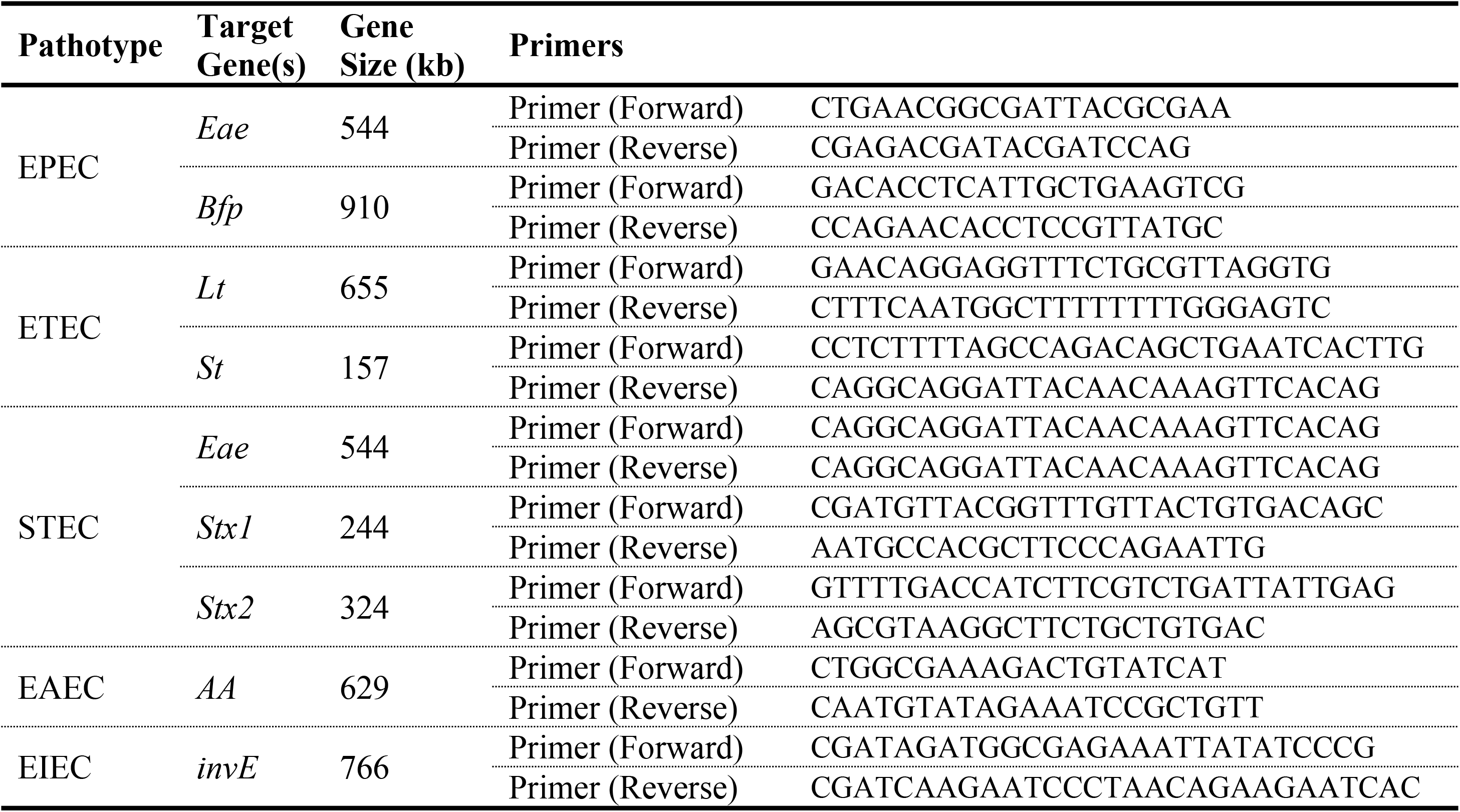
Target genes and their characteristics for isolation of different pathotypes.

For the preparation of samples, 10 *μl* Master Mix 2X (Fermentas, Catalog No. K0171) plus 7 *μl* of DDW and forward and reverse primers (1 *μl* of each) was added to 1 *μl* of sample. For positive and negative controls, 1 and kb DNA ladder and ladder mix were used. The processes of denaturation, annealing and extension were performed in the Eppendorf thermo cycler (Germany).

### Statistical analysis

Data were collected, revised, and analyzed by Stata software (version 14). Frequency of *E. coli* pathotypes was estimated in the overall population and in subgroups of age, location (province/city), and time (season/month). Frequency of *E. coli* pathogenic strains and target *E. coli* genes were estimated as the number of pathogenic *E. coli* samples identified by PCR tests to the total number of diarrhea samples with positive culture results for *E. coli* (presented as number and percent). Ninety-five percent confidence intervals for frequency measures were calculated with Wilson Score interval method. We grouped seasons into warmer (spring and summer) and cooler (fall and winter) seasons, between which the frequency of each *E. coli* pathotype was compared using the Chi Square test. Association of each *E. coli* pathotype with the type of diarrhea (with/without bleeding) was also assessed using the Chi Square test. Statistical tests were considered as significant at 0.05 levels.

## Results

In this study 1305 diarrheal samples were collected from 15 out of 31 provinces of Iran between January 2013 and January 2014. Of these, a total of 979 samples were *E. coli*-positive (75.0%; 95% CI: 72.6, 77.3%), and were subjected to molecular assays. Pathogenic *E. coli* strains were detected in 659 out of 1305 diarrheal samples, resulting in a prevalence of 50.5% (95% CI: 47.8, 53.2%).

STEC was the most and EIEC was the least frequent pathotypes among *E. coli*-positive diarrhea samples (frequency: 35.4% and 0.3%, respectively). The most frequent virulence genes identified included *stx1* and *Eae with* a frequency of 26.1% and 25.9%, respectively. The lowest frequent virulence genes included *bfp* and *invE*, each having a frequency of less than 1% (Table 2).

**Table 2.** prevalence of five *E. coli* pathotypes and their virulence genes in Iran.

| Pathotype | n | Prevalence in diarrheal samples* | Frequency in *E. coli*-positive samples** |
| --- | --- | --- | --- |
|  |  | % (95% CI) | % (95% CI) |
| **STEC** | 347 | 26.6 (24.2, 29.1) | 35.4 (32.4, 38.5) |
| **ETEC** | 137 | 10.5 (8.9, 12.3) | 14.0 (11.9, 16.3) |
| **EPEC** | 129 | 9.9 (8.3, 11.6) | 13.2 (11.1, 15.5) |
| **EAEC** | 43 | 3.3 (2.4, 4.4) | 4.4 (3.2, 5.9) |
| **EIEC** | 3 | 0.2 (0.1, 0.7) | 0.3 (0.001, 0.9) |
| **Total** | 659 | 50.5 (47.7, 53.2) | 67.3 (64.3, 70.3) |
| **Virulence genes** |  |  |  |
| ***Stx1*** | 255 | 19.5 (17.4, 21.8) | 26.1 (23.3, 28.9) |
| ***Eae*** | 254 | 19.5 (17.3, 21.7) | 25.9 (23.2, 28.8) |
| ***Stx2*** | 201 | 15.4 (13.5, 17.5) | 20.5 (18.0, 23.2) |
| ***LT*** | 80 | 6.1 (4.9, 07.6) | 8.2 (6.5, 10.1) |
| ***ST*** | 78 | 6.0 (4.8, 7.4) | 8.0 (6.5, 9.8) |
| ***AA*** | 43 | 3.3 (2.4, 4.4) | 4.4 (3.2, 5.9) |
| ***Bfp*** | 5 | 0.4 (0.1, 0.9) | 0.5 (0.2, 1.2) |
| ***invE*** | 4 | 0.3 (0.1, 0.8) | 0.4 (0.1, 1.0) |

\* Prevalence is calculated by dividing the numbers to the total number of diarrheal cases sampled (n= 1305). \*\*
Frequency is calculated by dividing the numbers to the total number of *E. coli*-positive samples identified in culture (n= 979)

### Geographical distribution of pathogenic *E. coli* in Iran

Pathogenic *E. coli* were detected in all investigated provinces. In this regard, frequency of *E. coli* pathotypes was highest in Fars province (S), where 88.7% of received *E. coli* samples were pathogenic (95% CI: 78.5, 94.4%). On the other hand, Razavi Khorasan Province (NE) had the lowest frequency among investigated provinces, although the prevalence in this province was still considerably high (34.8%; 95% CI: 35.5, 71.2%; Fig 1, Panel A, and S1 File, Table S1). At the city level, 100% of the received samples from Gorgan (NE; with 3 received sample), Tarem (WN; with 3 received sample), and Abadeh (S; with 9 received sample) were pathogenic *E. coli*. Pathogenic *E. coli* were not detected in three out of frothy-three cities. These cities included Saravan (SE; with 2 received samples), Damghan (CN; with 1 received sample), and Najaf Abad (C; with 5 samples; S2 File). Geographic distribution of five *E. coli* pathotypes is described below.

**Fig 1.**
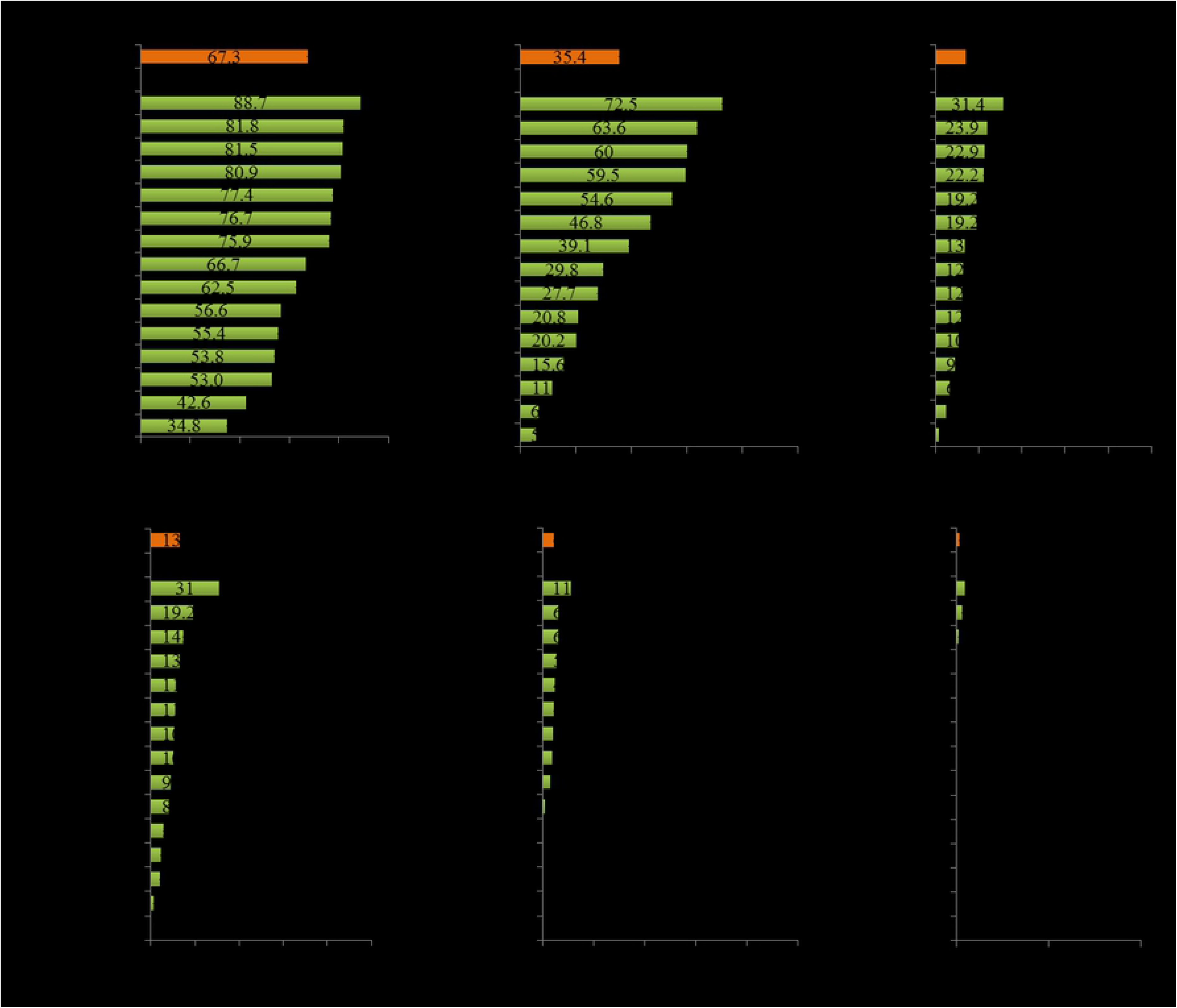
Frequency of *E. coli* pathotypes in 979 *E. coli*-positive culture samples collected in 15 provinces of Iran

**STEC**: was detected in all investigated provinces (Fig 1, Panel B). With an estimated frequency of 35.4% (95% CI: 32.4, 38.5%), STEC was identified as the most frequent pathotype. STEC constituted 52.7% of all identified pathogenic *E. coli* (S1 File, Table S2). The highest frequency of STEC was seen in Fars province (S; 72.5%). On the other hand, Esfahan had the lowest frequency of STEC (C; 5.5%; Fig 1, Panel B and S1 File, Table S2). At the city-level, STEC was detected in 33 out of 43 sampled cities (76.7%). In the majority of these cities (n= 21; 63.6%), STEC was also the predominant pathotype (Fig 2 panel A, and S2 File).

**Fig 2.**
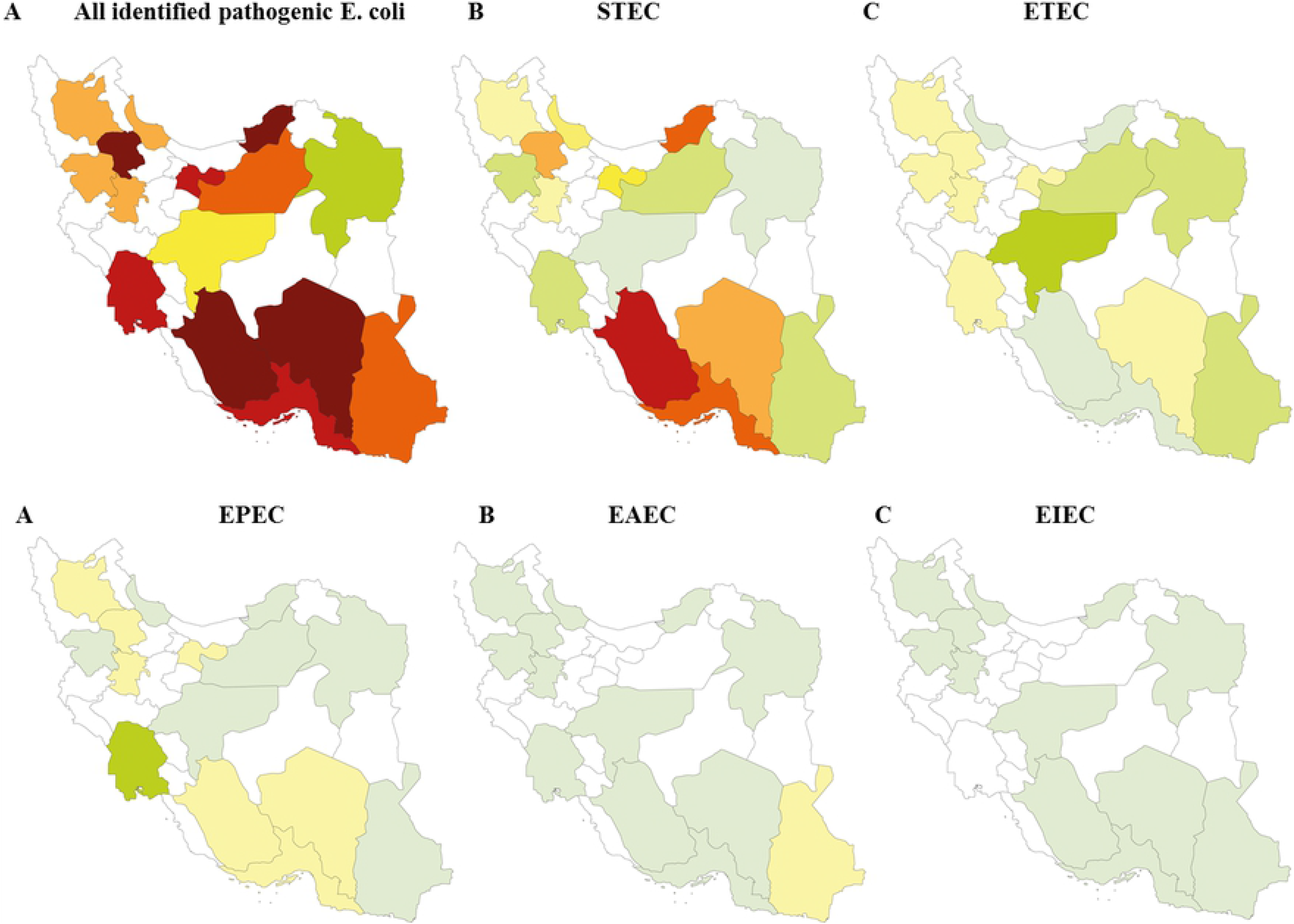
Distribution of five pathogenic *E. coli* in 15 selected provinces of Iran. The maps are originally developed by the authors of this manuscript.

**ETEC**: was also detected in all investigated provinces, although the number of identified ETEC isolates was quite lower than STEC (Fig 1, Panel C). With an estimated frequency of 14.0% (95% CI: 11.9, 16.3%), ETEC was identified as the second most frequent pathotype. ETEC constituted 20.8% of all identified pathogenic *E. coli* (S1 File, Table S3). At the provincial level, Esfahan province (C) had the highest ETEC frequency (31.4%), while Fars province (S) had the lowest frequency (1.6%) for this pathotype (S1 File, Table S3 and Fig 1, Panel C). At the city-level, ETEC was detected in 29 out of 43 cities (67.4%), and was the predominant pathotype in 9 of them (Fig 2 Panel B, and S2 File).

**EPEC**: was the third most common *E. coli* pathotype among all identified *E. coli* species in Iran, with a frequency of 13.1% (95% CI: 11.1, 15.4%). EPEC strains also constituted 19.6% of all identified ‘pathogenic *E. coli*’ (S1 File, Table S4). The highest frequency of EPEC (31.0%) was observed in Khuzestan province (SW). This pathotype was not observed in Esfahan province (C). Its frequency among *E. coli*-positive samples was also less than 5% in three provinces, including Razavi Khorasan (NE; 4.3%), Gilan (N; 4.0%), and Semnan (CN; 1.2%; S1 File, Table S4 and Fig 1, Panel D). At the city-level, EPEC was detected in 28 out of 43 cities (65.1%), and was the predominant pathotype in 4 of them (17.4%; Fig 2, Panel C, and S2 File).

**EAEC**: was not highly prevalent in Iran (4.3%; 95% CI: 3.1, 5.8%). This pathotype also constituted a small proportion of all identified pathogenic *E. coli* (6.5%; S1 File, Table S5). Except Kurdistan province (W) which had moderate number of positive EAEC pathotypes (n= 16), all other provinces identified few EAEC isolates. In fact, nine out of fifteen provinces identified ≤5 EAEC isolates, and five provinces did not identify this pathotype at all (S1 File, Table S5 and Fig 1, Panel E). At the city level, EAEC was detected in 19 out of 43 cities (44.2%). However, in most of these cities (n= 13) only one EAEC isolate was detected (Fig 2, Panel D, and S2 File).

**EIEC**: was a rare pathotype in Iran (Fig 1, Panel F). Only three EIEC strains were identified in our sample, which yielded an overall frequency of 0.3% (95% CI: 0.1, 0.9%; S1 File, Table S6). The three positive strains were detected in Sirjan (SE; 0.9%), Andimeshk (SW; 0.6%), and Semnan (CN; 0.2%) cities (Fig 2, Panel E, and S2 File).

### Temporal variation in the prevalence of *E. coli* pathotypes

STEC was more prevalent during summer and fall, and ETEC usually showed a pick during spring and summer. Notifiable seasonal trends could not be observed for other pathotypes, probably due to very few samples (Fig 3, Panel A). Analysis of monthly trends showed a pick for STEC frequency in March and a pick for ETEC frequency in June. Notifiable monthly trends could not be identified for other pathotypes, probably due to very few samples (Fig 3, Panel B). Our results also showed that *E. coli* pathotypes were generally more frequent in warmer (*i.e.*, spring and summer) than cooler seasons (*i.e.*, fall and winter). However, the difference was not statistically significant (69.1 vs. 65.9%, *P* value= 0.321). This seasonal pattern was observed for the ETEC pathotype and the seasonal trend was also statistically significant (17.6 vs. 11.3%, *P* value= 0.047). The same pattern was observed for EAEC and EIEC pathotypes, as well. However, the observed seasonal difference was negligible in size and was not statistically significant for these pathotypes. A reverse seasonal pattern was observed for STEC and EPEC, in a way that they were more frequently detected in cooler seasons than warmer seasons. The difference, however, was small in size and was not statistically significant (Fig 3, Panel C). The S3 File provides further details on the seasonal trend of five pathotypes in different provinces investigated in this study.

**Fig 3.**
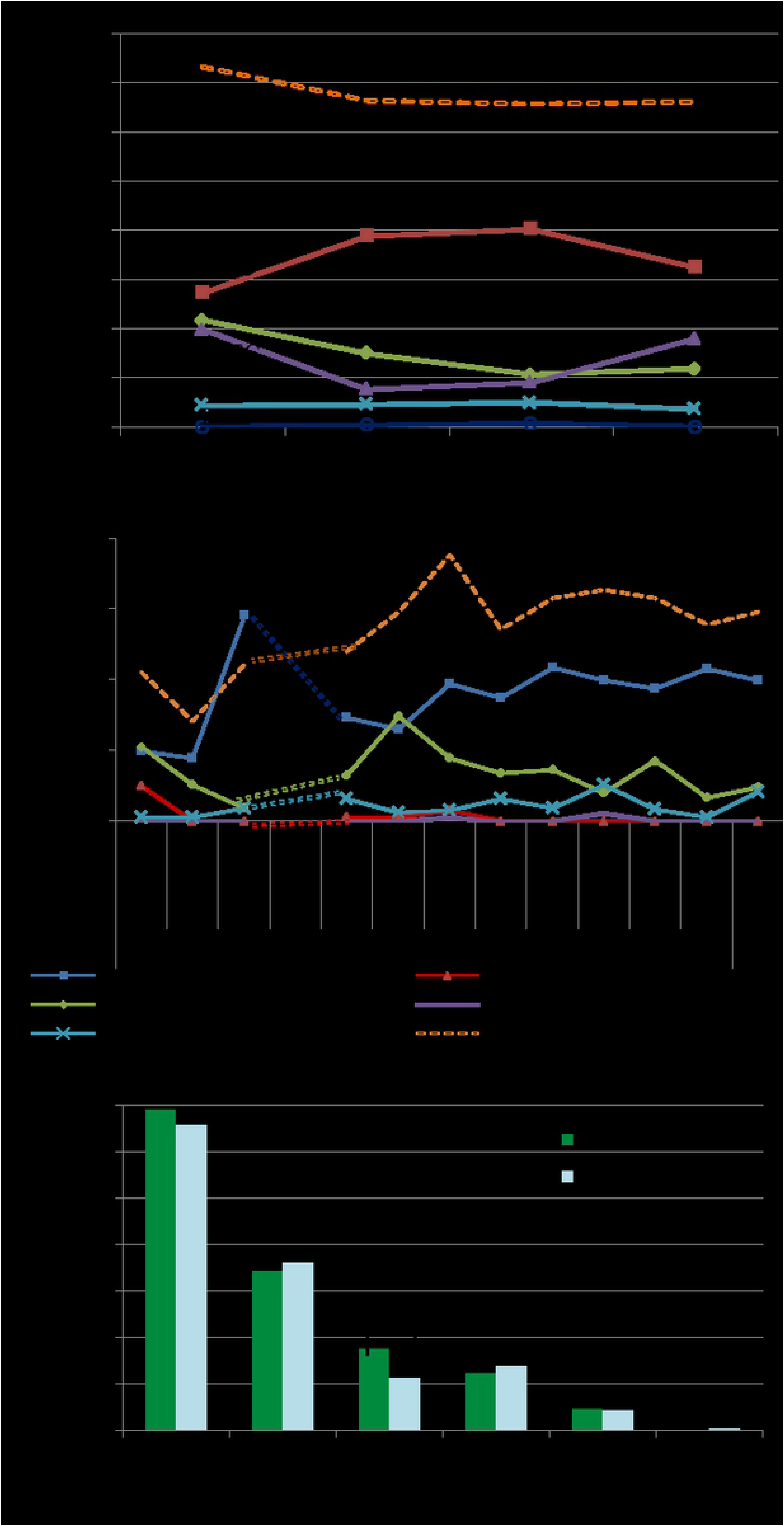
Seasonal trend of *E. coli* pathotypes in Iran. **(a) Frequency of five *E. coli* pathotypes in four seasons.** STEC was more prevalent during summer and fall, and ETEC usually showed a pick during spring and summer. Notifiable seasonal trends could not be observed for other pathotypes, probably due to very few samples. **(B) Frequency of five *E. coli* pathotypes in different months of a year.** STEC showed a pick in March and ETEC showed a pick in June. Notifiable monthly trends could not be observed for other pathotypes, probably due to very few samples. **(C) Difference in the frequency of five *E. coli* pathotypes in warmer (spring and summer) versus cooler seasons (fall and winter).** In overall, *E. coli* pathotypes were more frequently detected in warmer than cooler seasons but the difference was not statistically significant (*P* value= 0.321). This pattern was observed for the ETEC and EAEC pathotypes and was statistically significant for ETEC (*P* value= 0.04). STEC and EPEC were more frequently detected in cooler than warmer seasons but the difference, was negligible and not statistically significant.

### Age-specific prevalence of *E. coli* pathotypes

The highest frequency of pathogenic *E. coli* infections was observed in infants and children of 1-5 years of age (73% each). Most of the five *E. coli* pathotypes also had a high frequency among children. In this regard, STEC (40.5% and 41.1%, respectively) and EIEC (2.7% and 0.4%, respectively) pathotypes had the highest frequency among infants and children of 1-5 years. EPEC was highly frequent in children of 1-5 years, as well as in adults of 30-40 years (16.7% each). ETEC had the highest frequency in children of 5-10 years (21.4%). Although the frequency of pathogenic *E. coli* was higher in children, these pathotypes was also common in diarrhea samples received from adults and geriatrics (Table 3).

**Table 3.**
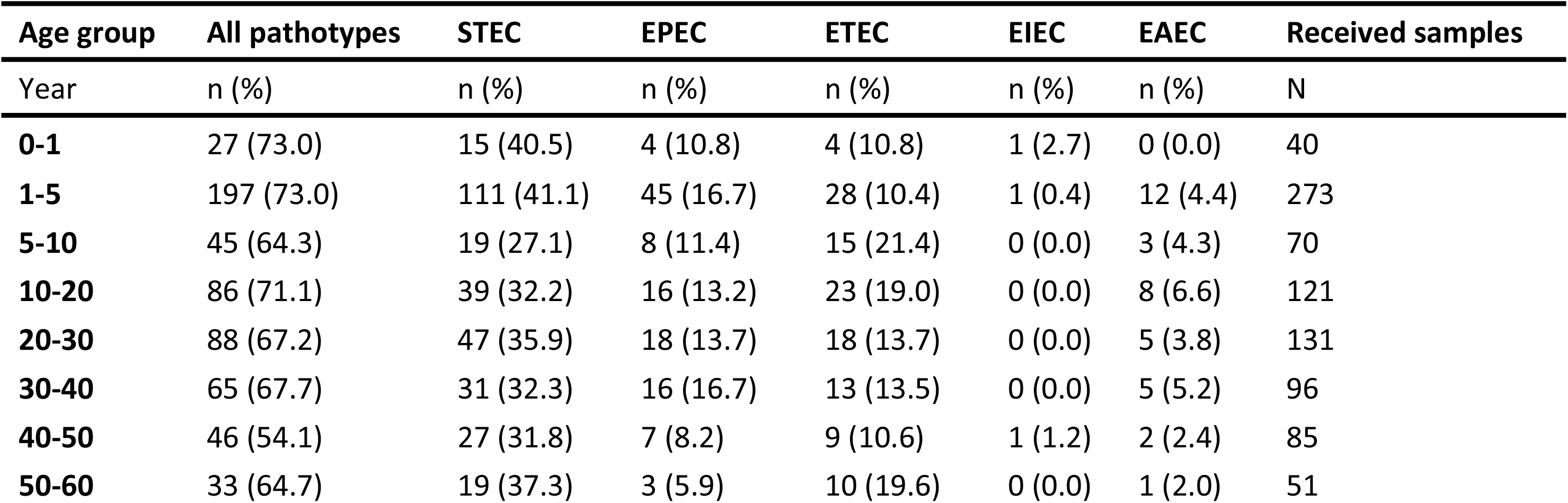
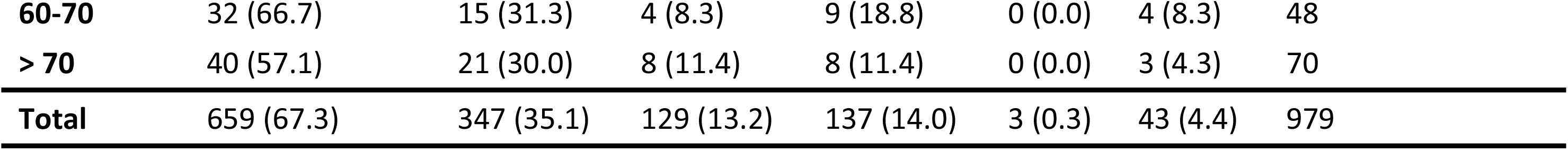
Age-specific frequency of *E. coli* pathotypes in 979 *E. coli*-positive culture samples received from 15 provinces of Iran

### Association with Sex

Our results showed no association between sex and infection with pathogenic *E. coli* pathotypes (data not shown).

## Discussion

In this large, nationally-representative survey in Iran, it was estimated that 31.6% of acute diarrhea cases were culture-positive for *E. coli*, among which 67.3% were infected with an *E. coli* pathotype. The study has yielded several findings about the frequency and spatial and temporal variation of *E. coli* pathotypes among cases with acute diarrhea in Iran.

STEC was the predominant pathotype in our sample, both at the national and provincial levels. STEC was also the most prevalent pathotype in infants and children under five years of age. Historically, STEC is among one of the first *E. coli* pathotypes isolated in Iran. The first report of this pathotype in Iran was published in 1998, where 2008 fecal samples were randomly selected from the general population of Ilam province (western Iran), and resulted in a prevalence of 4.9% for this pathotype (23). In a similar study conducted a few years later in the general population of two Northern provinces of Iran (Golestan and Mazandaran), the prevalence was estimated at 0.7%, which was quite lower than that reported from Ilam province (24). Both studies used cytotoxic methods for identification of STEC pathotypes in the Vero cells. The heterogeneity observed in the prevalence of STEC in these studies is consistent with our results, as we observed a considerable amount of heterogeneity in DEC frequency between investigated provinces (see Fig 1 and 2). Regional heterogeneity in the distribution of VTEC have been reported in other countries as well (25, 26). It has been well acknowledged that STEC pathotype has a strong negative association with age, with children under the five years of age being more susceptible to the infection. Therefore, the heterogeneity of STEC prevalence observed between 15 provinces investigated in this study could be attributed to heterogeneous age distribution in samples collected from these provinces.

Frequency of STEC pathotype in diarrheal children under five years of age was estimated at 7.8%, which is quite lower than similar estimates available from Iran (27–31). Molecular tests were used as the diagnostic method in all these studies. These results suggest that factors other than age also predict the extent of *E. coli* pathotype infection in the communities. In a study in Brazil that hierarchically analyzed data from 3725 children less than five years of age, Vasconcelos, et al. reported that environmental factors associated with the geographical area of residence, number of people per room, maternal age, and the age of the child are the factors that independently associate with diarrhea in the children (32). A similar study conducted later in Kenya (2013), reported that hand hygiene of the care-givers and the child, drinking untreated water from the river and lack of exclusive breastfeeding were significant predictors of child diarrhea (33). These factors would not be homogenous across provinces of Iran, and may justify the observed heterogeneity between different provinces and difference studies in the country.

ETEC is more prevalent in developing countries and is a major pathogenic strain in travelers’ diarrhea. This is consistent to our results, because ETEC was the second most prevalent pathotype in our study. This pathotype, however, had a moderate prevalence among children less than five years of age who participated in our study. Its prevalence was highest among adolescents and adults. Isolation and detection of ETEC is more difficult than other DEC strains, which makes it more difficult to isolate and detect it in the fecal samples of children and infants with diarrhea (16).

EPEC was the third most frequent pathotype in our study. It was also highly frequent in children under five years of age. EPEC is known to be among major causes of diarrhea in children, especially those under two years of age, with a prevalence of about 5-10% (34). Prevalence of EPEC in Iranian children has been reported by few studies. In two studies conducted in 2001 and 2006 on diarrheal children who were less than five years of age and referred to the community health centers of Eslam-Shahr (a suburban area of Tehran), EPEC prevalence was reported around 7% (35, 36). The same estimate was also reported by Salmanzadeh, et al (2005) in diarrhea children of Tehran (37). Two other studies conducted on diarrheal children of the same age who referred to four hospitals of Iran, reported an EPEC prevalence of 12.6 and 23% (28, 29). Higher prevalence of EPEC observed in the two latter studies, which selected hospitalized diarrheal children, might be indicative of the role of EPEC pathotype in severe diarrhea diseases. Prevalence of EPEC among Iranian adults has been less regarded in the literature. The very few available studies have focused on serological detection of EPEC. For example, Alikhani, et al. (2006 and 2007), reported an isolation rate of 44.9% and 36.4% for EPEC among diarrheal adults living in Ilam, Mazandaran, and Tehran provinces (38, 39). Since these estimates are based on serological tests, it might be impractical to compare them against our results which are based on PCR methods.

In our sample, EAEC was not frequently observed in adults and children. It constituted a small proportion of DEC pathotypes (about 6%). This pathotype has been recently acknowledged as a DEC, and is mainly regarded because of its role in travelers’ diarrhea. As we did not include travelers in our study, the lower frequency of EAEC in our sample can be justified. Very few studies have investigated the prevalence of this pathotype in Iran. The study conducted by Miri, et al. (2017) in Northern provinces of Iran, reported zero prevalence of EAEC in diarrheal patients of all ages who referred to community outpatients (40). These results are in line with our findings. Another study which was conducted by Jafari, et al. (2009), reported an EAEC prevalence of 16.3% in acute diarrhea children who were less than five years of age and hospitalized in children’s hospitals of Tehran (29). The rate is considerably higher than our estimates, and suggests that the prevalence of EAEC is higher in severe cases of diarrhea that lead to hospitalization. The role of EPEC and EAEC in severe diarrhea, especially among young children, would be an area that warrants further investigation.

There is no report on the prevalence and distribution of EIEC in Iran. As the first study in Iran, we found a frequency of 0.3% for this pathotype. Two out of three identified EIEC isolates identified in our study were detected in children less than five years of age. Human is the sole reservoir for the EIEC pathotype. Its transmission also requires high load of the pathogen, which decreases the chance of human-to-human transmission. Given these features of the pathotype and the very low prevalence of this pathotype in our study (which possesses a representative sample from Iran); it seems that infection with EIEC is a less important issue in Iran.

Our results showed that the overall frequency of pathogenic *E. coli* was higher in warmer seasons than cooler seasons. This was also the case for the ETEC and EAEC pathotypes, but a reverse pattern was observed for STEC and EPEC, in a way that their prevalence was slightly lower during warmer seasons than cooler seasons. These results are in line with previous studies in Mexico (41) and Kenya (42). These epidemiological findings could impact the recommended use of *E. coli* vaccines during warmer and cooler months. However, additional studies using pathotype-specific vaccines would be needed to further illuminate the possible benefits during lower acquisition rate seasons. The difference between ETEC-EAEC and STEC-EPEC rates in terms of seasonality suggests that the two pathotype groups have different pathways of transmission and reservoirs in Iran.

This study has notable strengths. Most studies on this topic in Iran are limited to one or two DEC pathotypes, in a specific age group and particular geographical areas. To the best of our knowledge, this is the first study that provides a nationally representative sample of diarrhea cases, which also includes all age groups and five major DEC pathotypes including STEC, ETEC, EPEC, EAEC, and EIEC. Half of the provinces in Iran were included in this study leading to a sampling proportion of 1:2 for provinces. We also sampled provinces in a way that an even distribution of sampled provinces is reached throughout the country. Our sampling also covered a time interval of 12 months, providing the opportunity to assess the rate of DEC pathotypes over a year. We hope that these attempts make the results more representative of the distribution and frequency of DEC pathotypes in the country.

This study has also a number of limitations that need to be considered. Due to financial constraints and large-scale nature of the study, we did not identify the serotype of isolated *E. coli*. Hence, the results represent the frequency estimates that are based on molecular detection of *E. coli* pathotypes. Further studies on the O, K, and H antigens of isolated pathotypes would provide a clearer picture of the distribution of major *E. coli* serotypes in the country. Antibiotic resistance profile of isolated pathogenic *E. coli* was not also investigated in this study. In our future work, we plan to determine the antibiotic resistance profile of *E. coli* pathotypes isolated in the current study. April coincides with the new year in Iran when most Iranians travel to other cities/countries. Therefore, diarrhea cases who refer to health centers during April are usually a mix of travelers and the local people. Given that we did not aim to study travelers’ diarhea in this study, we did not include diarheal cases referred to health centers during April. In order to provide a comprehension about DEC prevalence during April, we intrapolated the trend in this period. We also did not include hospitalized diarrhea cases as well as cases with chronic diarhea. So, the distribution of pathogenic agents responsible for these types of diarrhea in Iran remains unknown and warrants future research. Finally, existing prevalence studies in Iran are limited to few DEC strains, specific age groups, and limited geographical areas. The sampling method as well as pathogen isolation techniques are also heterogenous among these studies. These issues limited our ability to compare our results to those of previous studies. So, we could not derive conclusions about trends in the prevalence of DEC pathotypes. This issue highlights the need for development of similar national-level evaluations in the future. The results of such evaluations can then be used for the evaluation of time-trends and effectiveness of interventions and policies.

## Conclusion

The current study suggests that diarrheagenic *E. coli* may be an important cause of acute diarrhea both in adults and children in Iran, perhaps accounting for 7.2% and 10% of all acute diarrhea cases. Of five investigated pathotypes, STEC and ETEC seem to be widespread in Iran, which show a peak in warmer seasons (*i.e.*, spring and summer, respectively). This epidemiological finding could impact the recommended use of STEC and ETEC vaccines during warmer seasons, especially for infants, young children and elderlies. EPEC and EAEC seem to be less prevalent, and EIEC seems to be a rare pathotype in outpatients with acute diarrhea. The low EPEC and EAEC rates observed in our study and the high rate of these pathotypes reported among hospitalized children, suggests that the two important causes of diarrhea may be positively associated with severe diarrhea in children. Further investigations are warranted in this area. Finally, continued national evaluations of the prevalence of DEC pathotypes, as well as circulating strains and their antibiotic resistance profile, is recommended for evaluations of time-trends and effectiveness of interventions.

## Acknowledgements

The authors would like to thank the staff of health centers in selected provinces for their collaboration with this study. We also would like to thank laboratory staff at Molecular Biology Department in Pasteur Institute of Iran, for their important technical assistance.

## Supporting information

**S1 File. Frequency of *E. coli* pathotypes in 15 selected provinces of Iran**

**S2 File. Frequency of *E. coli* pathotypes in 43 selected cities of Iran**

**S3 File. Seasonal trend of *E. coli* pathotypes in 15 selected provinces of Iran**

